# Inferring monosynaptic connections from paired dendritic spine Ca^2+^ imaging and large-scale recording of extracellular spiking

**DOI:** 10.1101/2022.02.16.480643

**Authors:** Xiaohan Xue, Alessio Paolo Buccino, Sreedhar Saseendran Kumar, Andreas Hierlemann, Julian Bartram

## Abstract

Techniques to identify monosynaptic connections between neurons have been vital for neuroscience research, facilitating important advancements concerning network topology, synaptic plasticity, and synaptic integration, among others. Here, we introduce a novel approach to identify and monitor monosynaptic connections using high-resolution dendritic spine Ca^2+^ imaging combined with simultaneous large-scale recording of extracellular electrical activity by means of high-density microelectrode arrays (HD-MEAs). We introduce an easily adoptable analysis pipeline that associates the imaged spine with its presynaptic unit and test it on *in vitro* recordings. The method is further validated and optimized by simulating synaptically-evoked spine Ca^2+^ transients based on measured spike trains in order to obtain simulated ground-truth connections. The proposed approach offers unique advantages as *i*) it can be used to identify monosynaptic connections with an accurate localization of the synapse within the dendritic tree, *ii*) it provides precise information of presynaptic spiking, and *iii*) postsynaptic spine Ca^2+^ signals and, finally, iv) the non-invasive nature of the proposed method allows for long-term measurements. The analysis toolkit together with the rich data sets that were acquired are made publicly available for further exploration by the research community.

## 1 Introduction

Monosynaptic connections enable the flow of information between neurons. The contact point at which transmission occurs is the synapse, where the postsynaptic component of excitatory inputs is often a small protrusion called “dendritic spine”. The extent to which individual excitatory synapses contribute to the drive that the postsynaptic neuron receives depends on a multitude of synaptic properties, such as synaptic strength (with pre- and postsynaptic components), short-term plasticity processes, and the synapse location within the dendritic tree [1, 2, 3]. Moreover, the local dendritic environment, including adjacent synaptic inputs and dendritic properties, may allow for synaptic interactions and local dendritic information processing [4, 5]. The precise presynaptic spike patterns are, in this context, often of crucial importance. In addition, presynaptic activity and its relationship to postsynaptic spiking shapes the development of the synapse over time [6]. An in-depth interrogation of the functional properties, of organizational principles, and of the development of dendrites and their inputs requires, therefore, advanced methodologies, which provide both structural and functional information, ideally over extended periods of time.

Existing approaches for the identification and probing of monosynaptic connections have specific advantages and drawbacks [7, 8]. For example, axonal tracing in fixed tissue may allow for the exact localization of the synapse but cannot be used for functional investigations, while dual patch-clamp recordings grant, for a short period of time, access to pre- and postsynaptic activity, but the location of synaptic contacts is not directly available. Another approach to infer functional connectivity relies on correlated activity of extracellularly recorded spike trains [9, 10]. Here, the pairwise cross-correlation of unit spike trains is computed, based on which the spike-transmission probability (STP) between pre- and postsynatpic neuron can be assessed. While the non-invasive nature of this approach is certainly a strong advantage, STP values for a given connection depend on synaptic integration with other inputs and, consequently, they depend on the network state [11]. This implies that some connections may not be identifiable based on spike trains alone. Finally, methods that provide exquisite control over synaptic activation, either by electrical stimulation or using optical means (e.g., glutamate uncaging or optogenetics), do not provide access to precise spontaneous presynatpic spike times.

Here, we introduce a multimodal approach to map monosynaptic connections in functional neuronal networks that fills some of the existing gaps. To do so, we exploit the high temporal resolution of extracellular recordings and the high spatial resolution of Ca^2+^ imaging. Using simultaneous highdensity microelectrode array (HD-MEA) recordings and subcellular-resolution Ca^2+^ imaging in rat primary neuronal cultures, we were able to record precise spiking sequences of the many potential presynaptic cells and the single-spine Ca^2+^ transients of the evoked postsynaptic signals. We developed a novel and robust analysis pipeline to map monosynaptic connections on the basis of these paired recordings. We validate our method using data-driven simulations and demonstrate with several paired experimental recordings that the method can indeed be used to detect monosynaptic connections.

The presented approach represents a versatile tool for functional characterization of synaptic connections, including synaptic plasticity, activity-dependent synapse development and synaptic integration. In addition, location-dependencies of various synaptic properties can be probed.

All the data sets acquired in this work are available in the Neurodata Without Borders (NWB) format [12] on the DANDI archive https://gui.dandiarchive.org/#/dandiset/000223. The source code to implement the method and notebooks to reproduce the figures are available at: https://github.com/starquakes/mea_spineca_mapping.git.

## 2 Results

### 2.1 Identifying the presynaptic unit of a dendritic target spine

The proposed method for associating a specific dendritic spine with its putative presynaptic unit relies on a “brute-force” approach (Figure 1). We record the spiking activity of a large fraction of a neuronal network by means of high-density microelectrode arrays (HD-MEAs), while we simultaneously image the synaptically-evoked spine Ca^2+^ signals. Subsequently, we compute for each of the recorded spikesorted units an approximation of the spine Ca^2+^ trace that would be evoked in a postsynaptic neuron of the respective unit. Finally, a pairwise Pearson correlation between the measured spine Ca^2+^ transients and the computed traces is used to identify the most likely presynaptic unit.

**Figure 1:**
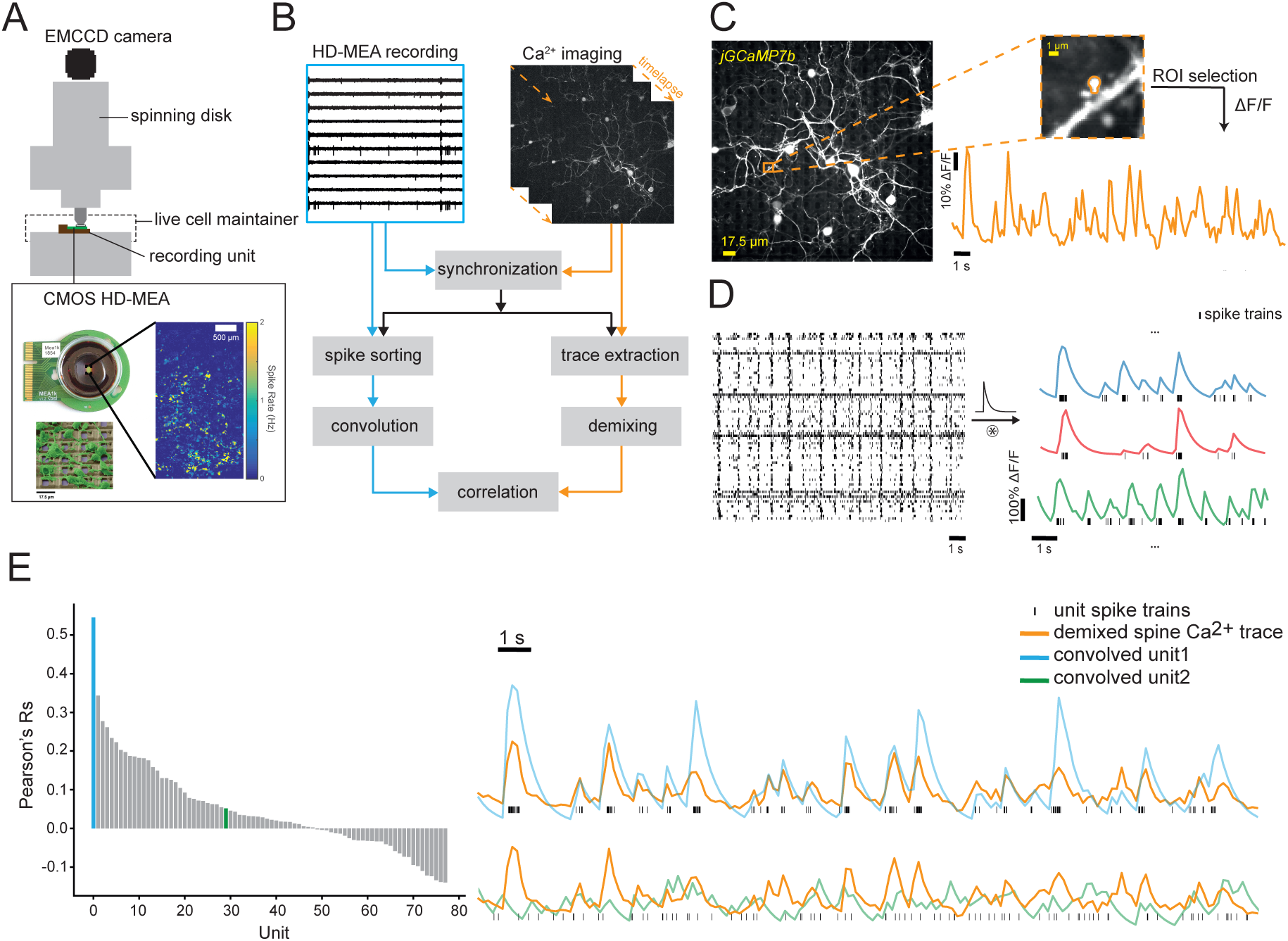
Experimental approach and analysis pipeline. **(A)** Upright confocal setup for simultaneously performing spine Ca^2+^ imaging and HD-MEA recordings. Sequentially recording from all 26’400 electrodes in blocks of 1’024 simultaneous recording channels makes it possible to generate a high-resolution spike-rate map of the entire network (inset, right; colors of some pixels are saturated for better overall visibility). **(B)** Overview of the analysis pipeline to match a spine Ca^2+^ trace with presynaptic activity. **(C)** Neurons with sparsely expressed jGCaMP7b plated on a HD-MEA chip (left). Magnified ROI containing a selected target spine, together with the extracted fluorescence trace of the spine (right). **(D)** Raster plot of spike-sorted HD-MEA data (left), and three example traces that are the result of convolving the respective spike trains with the depicted exponential decay kernel (right). **(E)** Sorted Pearson’s Rs (left) for one recording. Each R value is derived from the correlation of the measured and convolved unit with the spine Ca^2+^ trace. Comparison of the demixed measured spine trace (orange) with the convolved trace of the best-matching unit (blue) and a random unit (green) (right).

For data acquisition, an in-house-developed CMOS HD-MEA recording unit [13] with 26’400 electrodes at 17.5 μm pitch, featuring up to 1’024 channels that can be simultaneously recorded, was mounted under a spinning-disc upright confocal microscope (Figure 1A), as the HD-MEAs are not transparent. Primary rat cortical neurons were plated exclusively onto the active sensing area of the chip, and sparse expression of the fluorescent calcium indicator was achieved by co-transduction of floxed jGCaMP7b [14] and Cre with two adeno-associated viruses (AAVs). Experiments were performed at DIVs 18-21. The HD-MEA system allows for selecting an arbitrary configuration of the 1’024 electrodes that can be simultaneously used for recording, and different selection strategies are possible. Typically, our selection was based on a pre-scan of all 26’400 electrodes in order to determine electrodes that exhibit spiking activity (see Methods and Materials for details).

In Figure 1B, a more detailed overview of the processing pipeline is shown. Following temporal alignment, the HD-MEA and Ca^2+^ imaging data are further preprocessed in two separate streams. For Ca^2+^ imaging data, the first step requires a selection of the region of interest (ROI) containing the target spine (Figure 1C), followed by selection of an ROI containing the adjacent dendritic shaft (additional examples of Ca^2+^ indicator expression are shown in Figure 1 - figure supplement 1). The extracted Ca^2+^ traces are subsequently ΔF/F transformed. At this stage, the spine trace is a composition of Ca^2+^ signals associated with synaptic activations, but also backpropagating action potentials (bAPs). Since our method relies on correlations of synaptically-evoked signals, bAP-associated Ca^2+^ is an undesired component. Therefore, it is subsequently removed in a demixing step using the bAP-Ca^2+^-dominated shaft trace, as described before [15]. An example of the original ΔF/F of a spine and its adjacent dendritic shaft trace, as well as the demixed spine trace after removing the bAP-Ca^2+^ component, is shown in Figure 1 - figure supplement 2. The preprocessing of HD-MEA data includes two main steps: spike sorting and convolution. Spike sorting was performed using SpyKING Circus 0.8.4 [16] and SpikeInterface [17], followed by manual curation (using SpyKING Circus Matlab GUI). Next, we employed a decaying exponential kernel [18] that fitted the kinetics of the Ca^2+^ indicator [14] (decay *τ* = 0.5 *s*) to convolve each of the sorted spike trains (from now referred to as *convolved traces*). Finally, we downsampled the convolved traces to match the frame rate of the Ca^2+^ imaging. Examples of three convolved and downsampled traces are shown in Figure 1D.

**Figure 1 - figure supplement 1:**
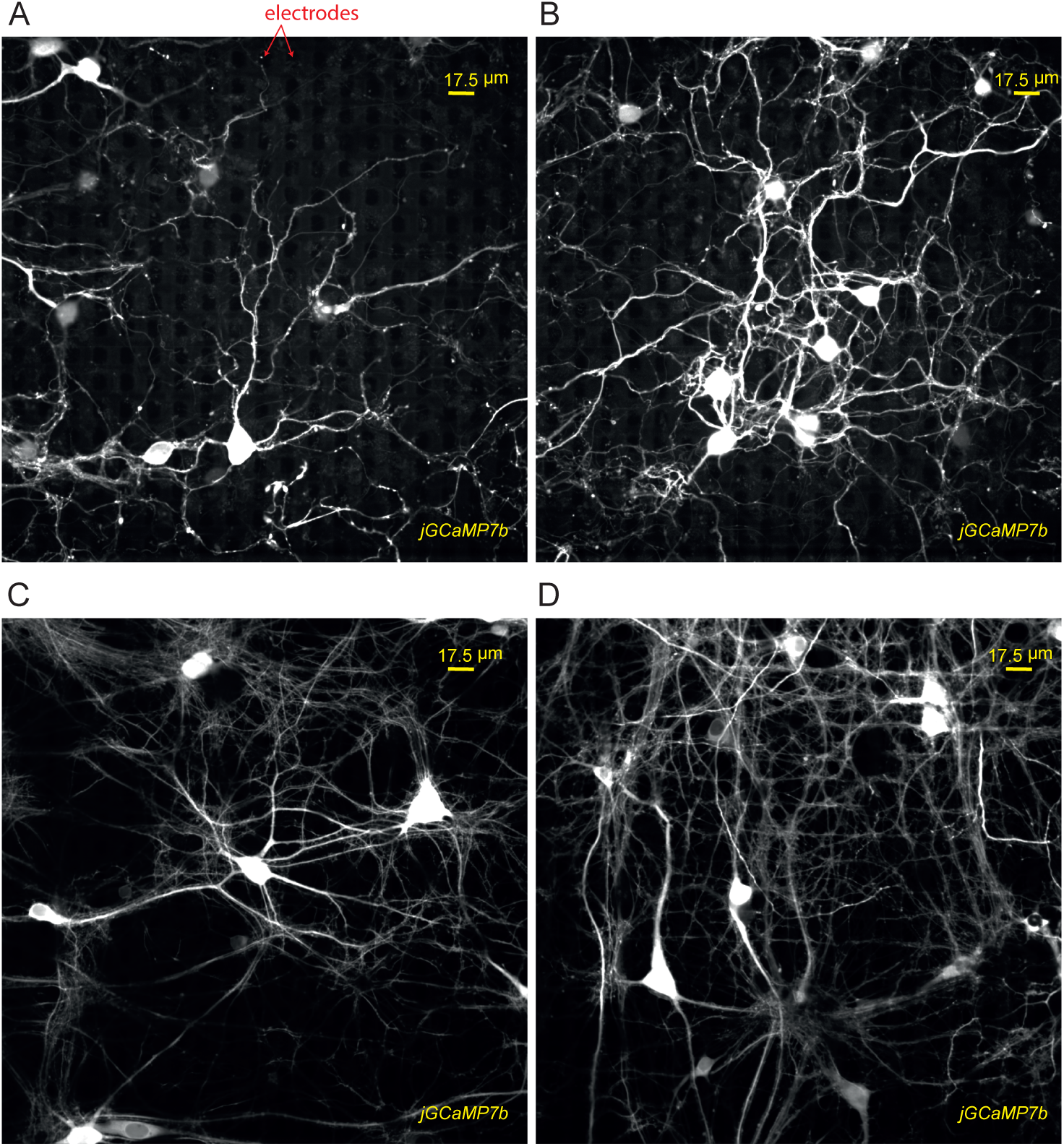
Four example Ca^2+^ imaging results, with neurons sparsely expressing jGCaMP7b plated on HD-MEA chips.

**Figure 1 - figure supplement 2:**
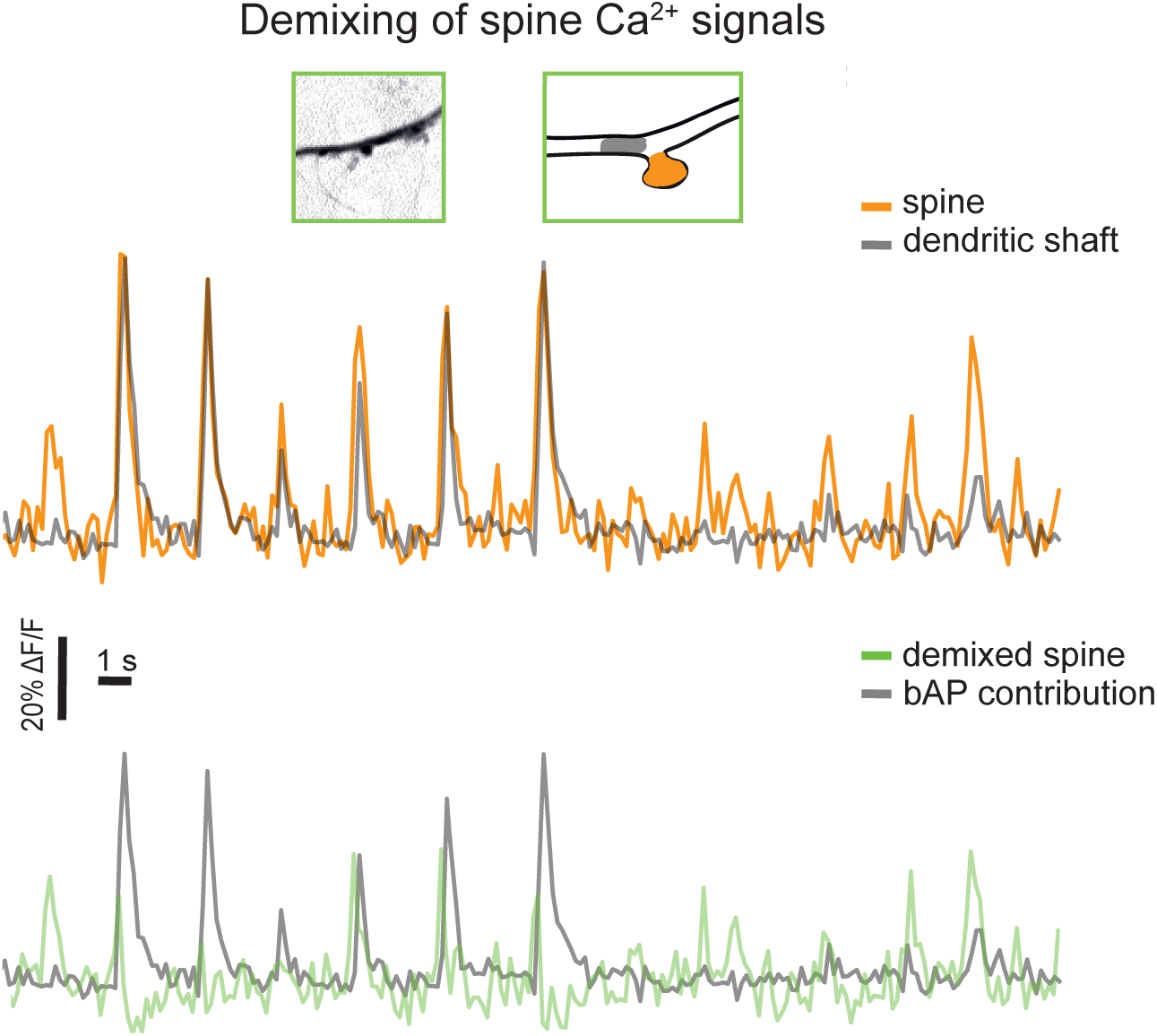
Demixing to isolate synaptically-evoked spine Ca^2+^ signals. The top panel shows the extracted ΔF/F traces of the spine (orange) and its adjacent dendritic shaft (grey). After demixing, individual spine activations (green) are isolated from bAP-associated Ca^2+^ signals and used for downstream analysis.

Since we aim to use correlations to infer synaptic connectivity, strong bursts of synchronous action potentials in the network will affect the possibility of identifying the presynaptic unit for a given spine. Moreover, increased non-linear postsynaptic interactions [19] during network bursting periods could introduce additional variation between measured and convolved spine Ca^2+^ traces. Therefore, an accurate detection and exclusion of network bursts is important for our analysis. By exploiting the network spiking information, we implemented a burst detection algorithm based on a global firing rate principle (see Methods and Materials) and excluded periods of strong bursting from the analysis (for an example, see Figure 1 - figure supplement 3).

As the final step to identify the presynaptic unit for the target spine, we computed the Pearson’s correlation between the demixed spine trace and the convolved traces from all recorded units (excluding the bursting periods). The recorded units could then be sorted by their Pearson correlation coefficient (Pearson’s R) (Figure 1E), and the best matching one (from now referred to as *best-matching unit*) was identified as the candidate to form a monosynaptic connection to the target spine. Note the remarkable correlation of presynaptic spike times and the convolved trace (blue) for the best-matching unit and the demixed measured spine Ca^2+^ trace (orange) - especially for smaller events. Moreover, the R value for the best-matching unit (*best-R*) shows the largest increase with respect to the next (ordered) R value among all units.

**Figure 1 - figure supplement 3:**
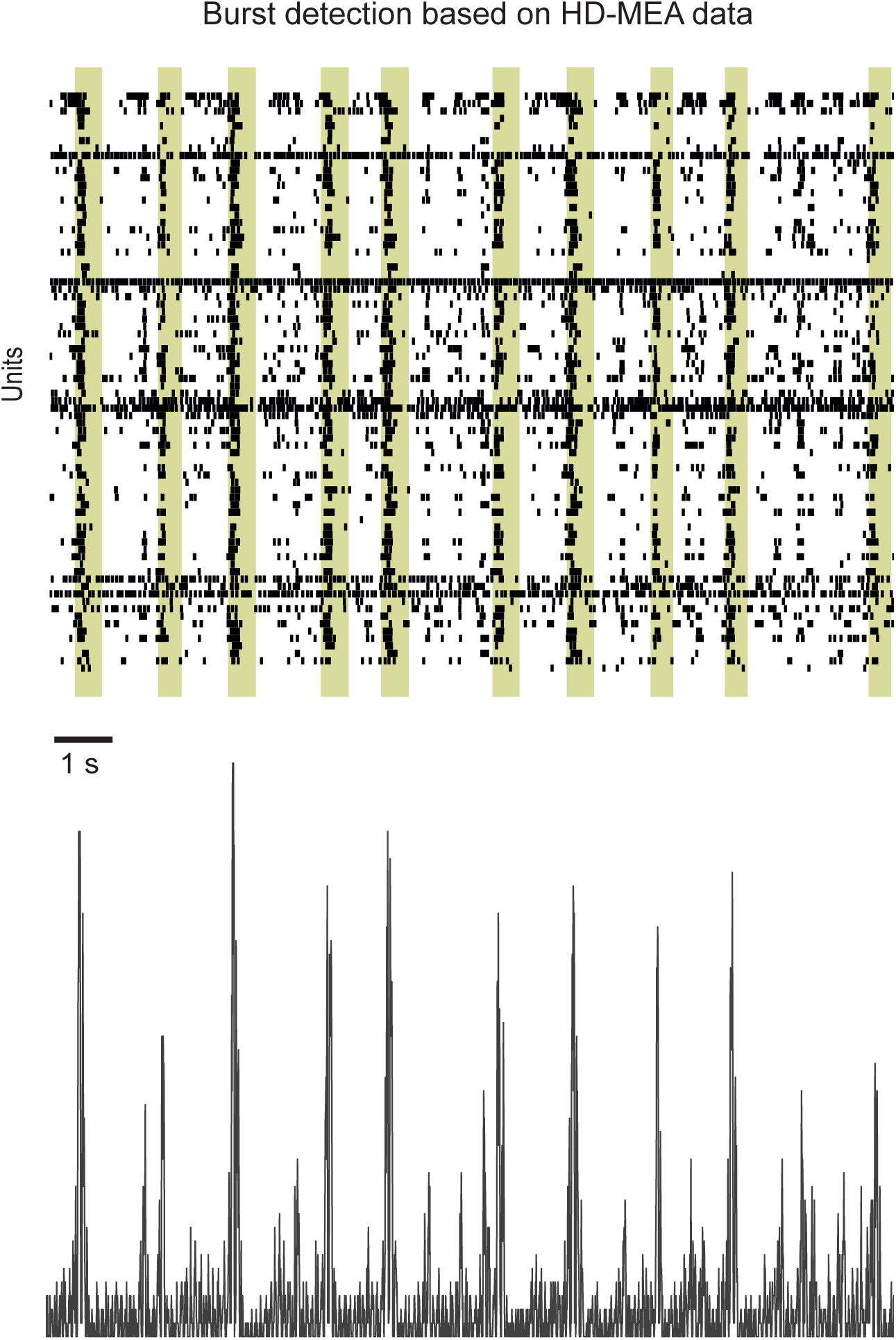
Network burst detection based on HD-MEA data. The top panel shows a raster plot of spike-sorted extracellular data recorded by the HD-MEA system. Each row shows the spike times of one unit. Detected network bursting periods are highlighted in yellow. The bottom panel shows the time-binned number of spikes of all units (bin size is 10 ms).

### 2.2 Acceptance test for putative monosynaptic connections

In the previous section, we outlined the experimental approach and analysis pipeline to identify a presynaptic candidate unit. However, *i*) an incomplete coverage of the network or *ii*) synchronous firing of neurons could result in a unit displaying the strongest correlation, although it is not the presynaptic neuron of the target spine. The latter is a possibility because presynaptic release probabilities and recording noise will cause stochastic deviations between measured demixed and convolved trace. Therefore, a more formal procedure to accept only a “trustworthy” presynaptic candidate unit and its putative connection is required. Here we introduce a novel data-driven approach, based on the surrogate method, to perform such an acceptance test.

We propose that each putative connection needs to meet two criteria to be accepted. These two criteria assure that only the spike train of a specific candidate presynaptic unit can, with reasonable confidence, be assumed to generate the measured spine Ca^2+^ transients. First, we require the best-R value to fall within a distribution of possible R values for the best-matching unit, assuming biologically plausible presynaptic release probabilities (P_r_) and realistic recording noise. To build such a distribution, we construct simulated ground-truth connections of the best-matching unit, i.e., we generate surrogate spine Ca^2+^ traces evoked by the best-matching unit spike train (here referred to as *ground-truth surrogates*; Figure 2).

**Figure 2:**
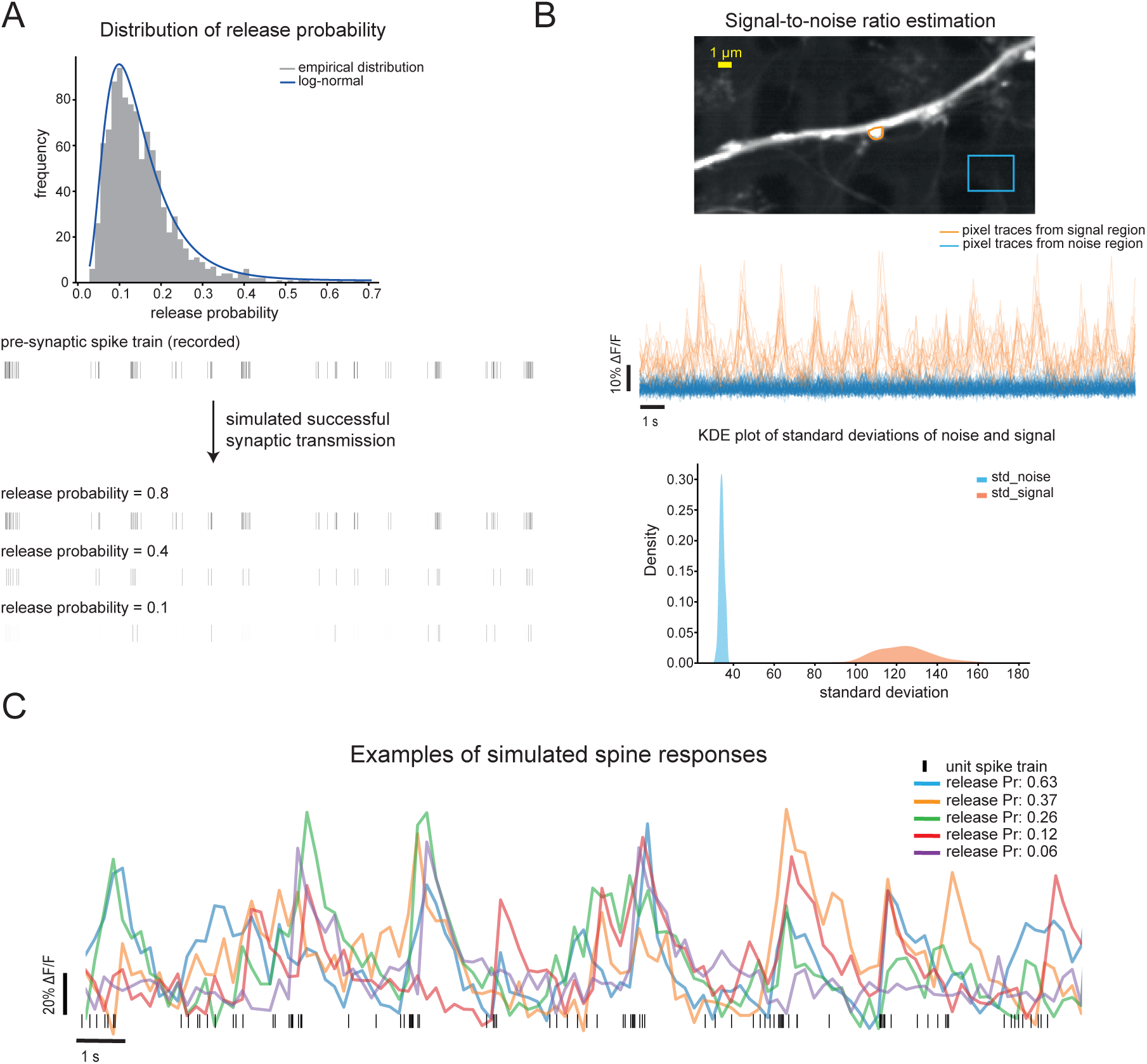
Surrogate method for the simulation of spine Ca^2+^ responses for a given presynaptic spike train. **(A)** Log-normal distribution of presynaptic release probabilities used for surrogate generation (top). We show three examples of simulated “successful synaptic transmissions”, based on different release probabilities, for the same measured presynaptic spike train (top). **(B)** Measuring the signal and noise levels of Ca^2+^ imaging data. Individual pixel traces were extracted from manually drawn ROIs containing the spine (signal) and a background region (noise) (top and middle). The kernel density estimation (KDE) plot of the standard deviations of the pixel traces of signal and noise is shown as well (bottom). **(C)** Examples of simulated surrogates of spine Ca^2+^ responses of the same spike train (black marks) incorporating different release probabilities and recording noise.

Essentially, this distribution can be obtained by, again, convolving the unit spike train with the exponential decay kernel that mimics the spine Ca^2+^ response. However, this time, we introduce two layers of uncertainty. The first one is given by the release probability of the synapse (Figure 2A). Here, we assume a log-normal distribution of P_r_ values that is consistent with previous reports [20]. For each surrogate generation for a given “presynaptic” spike train, a P_r_ value is drawn from this distribution and then determines the probability of the spikes that are retained as simulated successful synaptic transmissions. Several noise sources during Ca^2+^ microscopy introduce additional variations in the recorded spine traces. To account for these variations, we also estimate the signal-to-noise ratio for each target spine (Figure 2B) and add accordingly scaled white-noise to our surrogate traces (see Methods and Materials for details). Following these steps, an arbitrary number of spine Ca^2+^ trace surrogates can be generated for a given presynaptic spike train (Figure 2C). For our first acceptance criterion, we generated 1’000 spine trace surrogates based on the spike train of the best-matching unit and then calculated for each surrogate the Pearson’s R with the convolved trace of the best-matching unit (blue *ground-truth surrogate* data in Figure 3). In this way, we generated a distribution of possible Rs (*GT-Rs*) for our candidate presynaptic unit. We required the best-R value to fall “well within” the GT-R distribution, as will be further detailed below.

**Figure 3:**
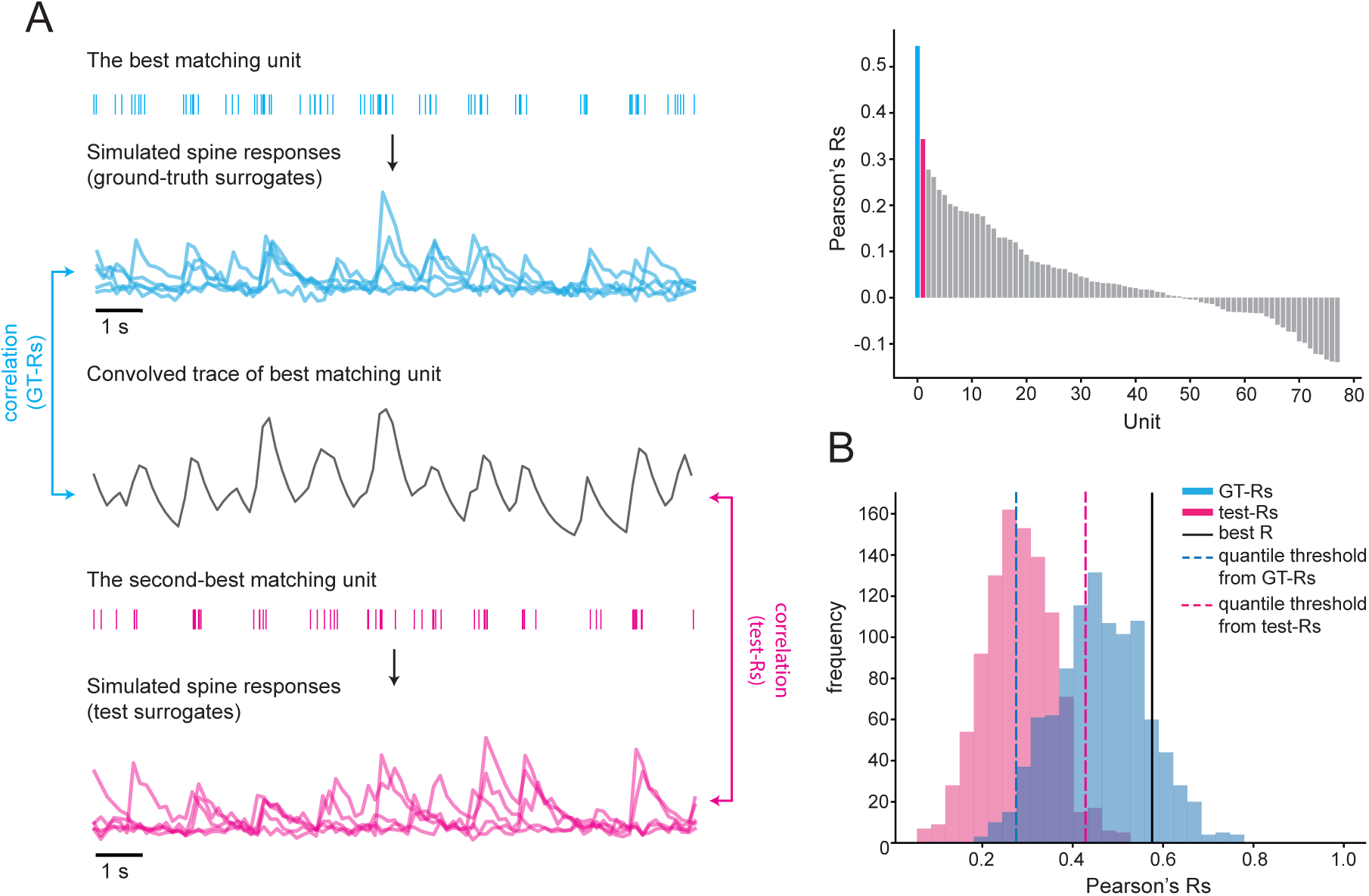
Acceptance test. **(A)** Simulated ground-truth surrogates based on the best-matching unit (blue) and test surrogates based on the second-best matching unit (magenta). **(B)** Distributions of GT-Rs and test-Rs are plotted together with the experimentally obtained best-R value. As an initial acceptance test, the 95^th^ quantile of the test-Rs (dashed magenta line) and 10^th^ quantile of the GT-Rs (dashed blue line) were selected to determine threshold values that the best-R value (black line) was required to exceed for a putative monosynaptic connection to be accepted.

This first criterion was introduced to assure that Ca^2+^ traces that could be evoked by the best-matching unit were compatible with our experimentally observed best-R value. However, to be able to *trust* this putative connection, we next needed to exclude that the best-R value also would be a likely outcome for any of the other units. This next step could be reduced to conducting a test for the unit with the second largest R (*second-best matching unit*) of the correlation between the demixed spine Ca^2+^ trace and convolved traces. Essentially, we asked here: Are the data compatible with the case that the second-best matching unit would be connected to our target spine? For this test, we again generated a population of 1’000 surrogate spine responses, but this time, we used the spike train of the second-best matching unit (magenta *test surrogate* data in Figure 3). Also for these surrogates, we subsequently calculated the distribution of Pearson’s R (*test-Rs*) by correlating each test surrogate with the convolved trace of the best-matching unit. Our second criterion for the acceptance of a putative monosynaptic connection was then that the best-R did not fall “well within” the test-Rs distribution. This means that the spike train of the second-best matching unit was unlikely to evoke spine Ca^2+^ responses that were compatible with our experimentally observed best-R value. Examples of test-R and GT-R distributions, together with the best-R values for this recording, are shown in Figure 3B.

With these distributions, we could now perform the acceptance test based on our two criteria. Finally, it remains to be formalized what is meant by the best-R falling (not) “well within” a distribution. Empirically, one could choose different quantiles in order to obtain the threshold values for the GT-R and test-R value distributions (for example, in Figure 3B we used the 95^th^ quantile for test-Rs and the 10^th^ quantile for GT-Rs). In the next section, we will show how to determine data-driven optimized quantile thresholds that maximize the method’s performance.

### 2.3 Method validation and threshold optimization

As part of the acceptance test, described in the previous section, we applied fixed quantiles (95^th^/10^th^) to the two R distributions (test-R/GT-R) in order to obtain thresholds for the best-R value. However, for best performance, these quantiles should be tailored to each network, where factors such as network synchrony may influence the optimal quantile thresholds.

To validate our method and to obtain the optimal quantile thresholds for the acceptance test, we further extended the surrogate method described in the previous section. For each recorded unit *i*, we generated *N* surrogate spine Ca^2+^ response traces. Each of these “presynaptic” units could then be considered to be part of a simulated ground-truth connection, and the surrogates constituted simulated spine Ca^2+^ measurements (Figure 4A). Next, we computed the GT-Rs and test-Rs distributions and we performed the acceptance test with a grid search over quantile values for the test-Rs (Q1) and the GT-Rs (Q2) (Figure 4B). From the outcome of the test over all surrogates and all units, we could compute the number of True Positives (TP), when unit *i* was accepted as a monosynaptic connection, and False Negatives (FN), when unit *i* was rejected as a monosynaptic connection (or another unit was found). In addition, we ran the test after removing the presynaptic unit *i*, based on which the surrogates were generated, from the list of recorded units. In this case, the test could yield True Negatives (TN), when the putative connection was rejected, or False Positive (FP), when a monosynaptic connection was accepted despite the fact that the presynaptic unit *i* was removed (Figure 4A). After counting all TPs, FNs, TNs, and FPs, we computed the *F*-score for each pair of Q1-Q2 as a performance metric. Figure 4C shows the heat map of *F*-scores for one recording, with the green arrow showing the maximum score and, therefore, the optimal thresholds for this specific network.

**Figure 4:**
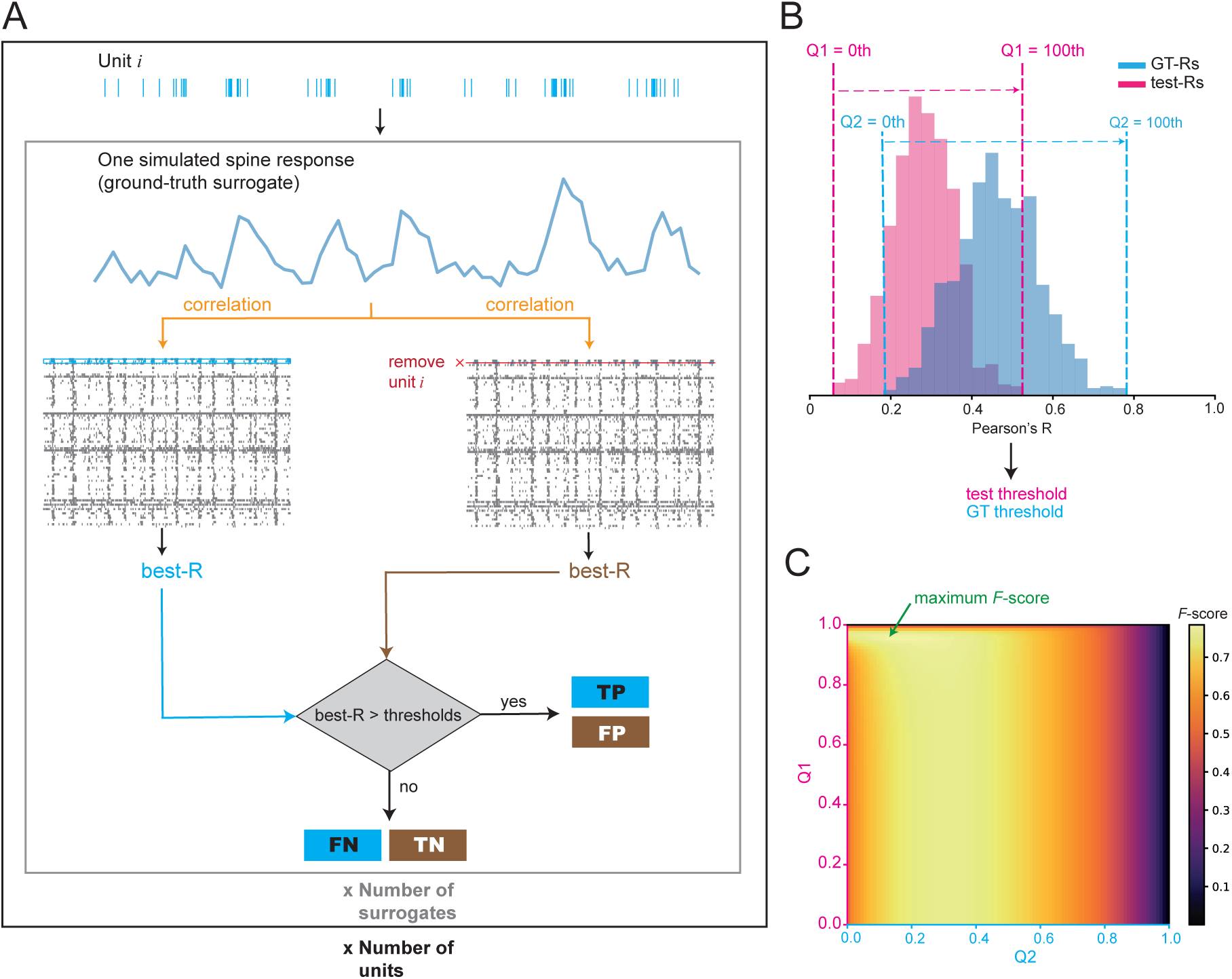
Method validation and threshold optimization. **(A)** Illustration of how ground-truth surrogates were used to calculate True Positives (TP), False Negatives (FN), True Negatives (TN) and False Positives (FP) to determine the *F*-score for performance evaluation. **(B)** Example GT-R and test-R distributions, with an indication of the range of values for the two quantiles Q1 and Q2. **(C)** Performance matrix for one example network. For each pair of Q1 and Q2, the algorithm, detailed in A, was executed to determine the corresponding *F*-score. The green arrow indicates the maximum of the *F*-score.

### 2.4 Performance evaluation on experimental data

To evaluate the performance of the proposed method upon application to real experimental data, we conducted 19 paired spine Ca^2+^ imaging and HD-MEA recording experiments in 10 different cell cultures. First, the optimal threshold values for the acceptance test were computed based on the spike-sorted unit data. From each recording, we then typically selected one medium-large dendritic spine that exhibited signs of activity (in one of recordings two spines were chosen). Table 1 shows the full analysis results. For 9 out of 20 spines, the best-matching unit passed the acceptance test, and a monosynaptic connection was confirmed. Moreover, the typically large *F*-score, associated with the optimal threshold values that were used, implied that the network characteristics (e.g. synchrony) permitted a good classification in most cultures.

**Table 1:** Results of tested monosynaptic connections.

| culture ID | spine ID | best_R | GT_Q | GT_thr | test_Q | test_thr | max_F | accept? | #units | OUT/IN ratio | spike contrast |
| --- | --- | --- | --- | --- | --- | --- | --- | --- | --- | --- | --- |
| 1 | 1 | 0.523 | 0.091 | 0.214 | 0.960 | 0.296 | 0.787 | yes | 78 | 0.733 | 0.38 |
|  | 2 | 0.516 | 0.162 | 0.291 | 0.960 | 0.271 | 0.854 | yes | 132 | 2.003 | 0.17 |
|  | 3 | 0.217 | 0.172 | 0.112 | 0.960 | 0.190 | 0.835 | yes | 66 | 1.717 | 0.17 |
|  | 4 | 0.644 | 0.162 | 0.311 | 0.949 | 0.489 | 0.850 | yes | 185 | 2.086 | 0.23 |
|  | 5 | 0.429 | 0.162 | 0.358 | 0.929 | 0.520 | 0.797 | no | 159 | 0.318 | 0.26 |
| 2 | 6 | 0.647 | 0.192 | 0.448 | 0.929 | 0.619 | 0.599 | yes | 50 | 0.189 | 0.41 |
|  | 7 | 0.346 | 0.152 | 0.503 | 0.929 | 0.585 | 0.604 | no | 55 | 0.269 | 0.48 |
|  | 8 | 0.340 | 0.030 | 0.227 | 0.929 | 0.475 | 0.544 | no | 66 | 0.060 | 0.62 |
| 3 | 9 | 0.671 | 0.222 | 0.507 | 0.949 | 0.639 | 0.795 | yes | 179 | 0.425 | 0.28 |
|  | 10 | 0.765 | 0.222 | 0.507 | 0.949 | 0.639 | 0.795 | yes | 179 | 0.425 | 0.28 |
|  | 11 | 0.148 | 0.232 | 0.354 | 0.939 | 0.351 | 0.775 | no | 99 | 0.783 | 0.20 |
| 4 | 12 | 0.597 | 0.152 | 0.353 | 0.960 | 0.544 | 0.870 | yes | 114 | 0.719 | 0.29 |
|  | 13 | 0.160 | 0.172 | 0.267 | 0.960 | 0.068 | 0.860 | no | 128 | 0.425 | 0.36 |
| 5 | 14 | 0.394 | 0.111 | 0.456 | 0.869 | 0.435 | 0.485 | no | 133 | 0.289 | 0.64 |
|  | 15 | 0.400 | 0.091 | 0.436 | 0.879 | 0.604 | 0.520 | no | 133 | 0.222 | 0.69 |
| 6 | 16 | 0.676 | 0.424 | 0.383 | 0.545 | 0.666 | 0.525 | yes | 58 | 0.171 | 0.43 |
| 7 | 17 | 0.602 | 0.232 | 0.500 | 0.889 | 0.707 | 0.537 | no | 50 | 0.534 | 0.31 |
| 8 | 18 | 0.311 | 0.182 | 0.214 | 0.949 | 0.324 | 0.723 | no | 100 | 0.464 | 0.25 |
| 9 | 19 | 0.219 | 0.182 | 0.233 | 0.909 | 0.298 | 0.610 | no | 87 | 0.272 | 0.50 |
| 10 | 20 | 0.294 | 0.121 | 0.414 | 0.879 | 0.364 | 0.577 | no | 59 | 0.112 | 0.53 |

We continued our investigation by probing in more detail the basis underlying the acceptance or rejection of a connection. Although the accepted connections had significantly larger best-R values (0.584 ± 0.149) compared to the rejected ones (0.331 ± 0.124) (Mann-Whitney *U* =12.0, *n*_1_=9, *n*_2_=11, *P* <0.05), it is important to note that the value of the best-R alone was not a predictor for the outcome of the test. In fact, as one can see from Figure 5B, the acceptance test rejected connections with R values as high as ~0.6, while in another case an R value of ~0.2 was enough for acceptance. Figure 5E shows two examples of accepted connections, while Figure 5F displays two rejected ones. Note that for some recordings the test threshold was larger than the GT threshold and vice versa for other recordings. Indeed, we found that putative connections were rejected in three different scenarios: The best-R was smaller than either one of the two thresholds or both thresholds (see summary of all tested spines in Figure 5A). This finding demonstrates the performance advantage of having two thresholds in the acceptance test.

**Figure 5:**
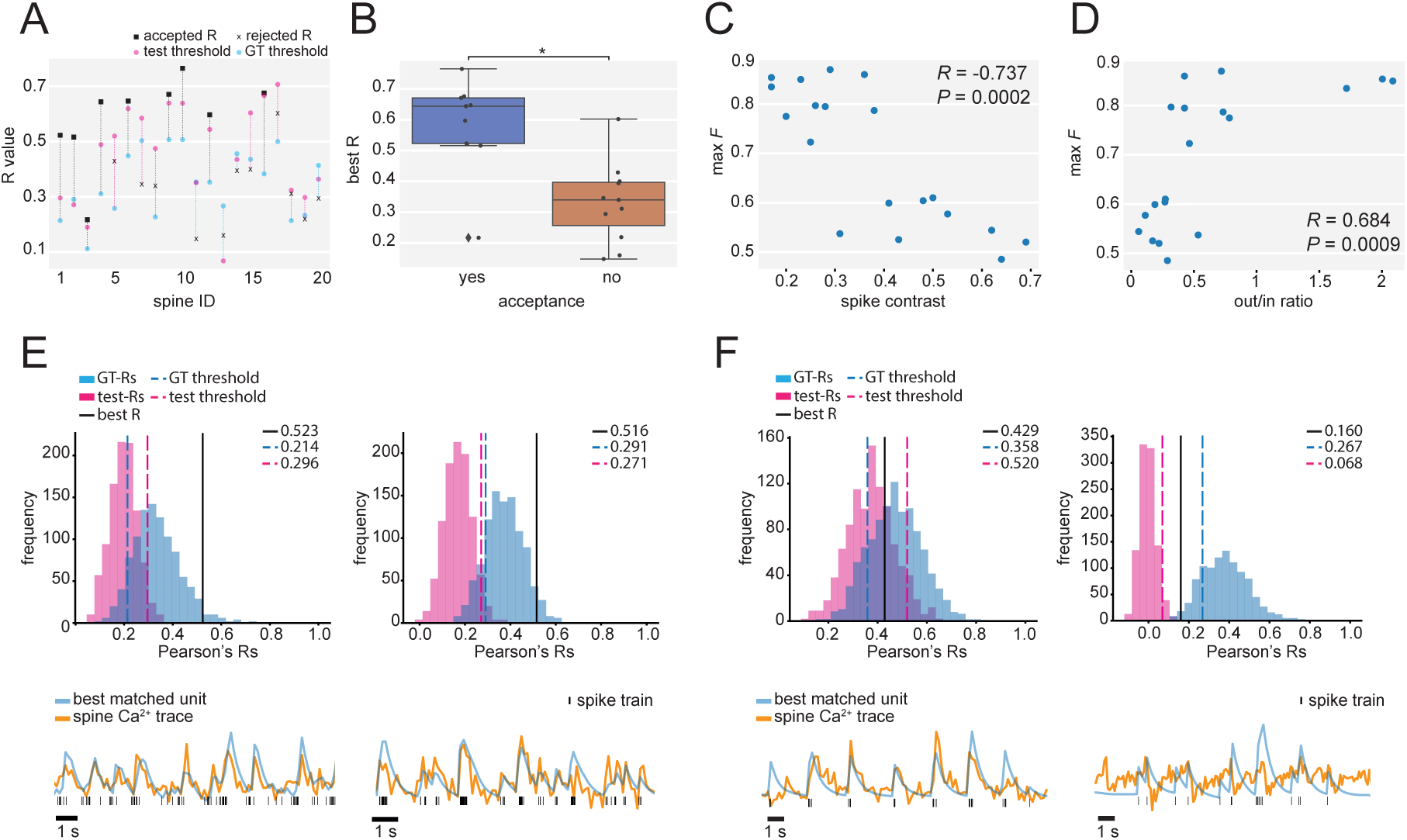
Application of connection inference method on experimental data. **(A)** Summary plot displaying, for each spine, the best-R (black squares or “x” for accepted and rejected connections, respectively) and the two threshold values based on the test-R (magenta circle) and GT-R (blue circle) distributions. Black dashed lines indicate accepted connections (best-R > two thresholds). Magenta or blue dashed lines indicate rejected cases based on the best-R not passing the test and GT threshold, respectively. **(B)** Box plot of best-Rs for accepted and rejected connections. **(C)** Maximum *F* score versus spike contrast. **(D)** Maximum *F* score versus ratio of spikes outside/inside bursts. **(E)** Two examples, in which the best-matching unit was accepted to form a monosynaptic connection with the imaged spine. Shown are the R distributions, thresholds and the best-R value for each connection (top). Furthermore, the measured spine Ca^2+^ trace and the convolved trace for the best-match unit are plotted at the bottom. **(F)** Two examples, in which the best-matching unit was rejected. Data displayed as in E. The reason for rejection varied: the experimental best-R was rejected by not passing the test threshold (left) or the GT threshold (right).

Finally, we further examined the hypothesis that network activity characteristics would affect the performance of our method. We calculated two complementary measures for each recording. First, we determined the ratio of spike events occurring outside and inside of network burst periods (OUT/IN column in Table 1), which captured to what extent neuron spiking was entrained by network bursting. Secondly, we calculated the spike contrast of the network, a fast measure of spike train synchrony [21] (implemented in the Elephant Python package [22] – version 0.10.0). We found that both the spike contrast (*R*=0.737, *P* =0.0002, Spearman’s rank correlation, Figure 5C) and the OUT/IN ratio (*R*=0.684, *P* =0.0009, Spearman’s rank correlation, Figure 5C) of a network correlated with the classification performance as measured by the maximum *F*-score of the respective network. We have, therefore, ways to screen for the most promising cultures, in terms of identifying monosynaptic connections, before even beginning with spine Ca^2+^ imaging.

## 3 Discussion

In this paper, we introduced a novel approach for mapping monosynaptic connections using subcellular resolution Ca^2+^ imaging of dendritic spines and simultaneously recorded extracellular electrical spiking activity from large-scale networks by means of HD-MEAs. We established a data-driven analysis pipeline that is able to match an optically recorded spine trace with the spiking activity of the putative presynaptic unit and accepts or rejects the connection based on a surrogate strategy. We validated and assessed the performance of the proposed method using simulated ground-truth monosynaptic connections and showed how the developed method can provide a robust detection of monosynaptic connections in experimental data. Our evaluation was performed with 2D-cell-culture recordings. Indeed, the possibility to control the number of cells in the network and the fact that HD-MEA systems can simultaneously record from large fractions of the grown networks, renders dissociated 2D cultures, plated on the HD-MEA chips, particularly suitable for the developed connection mapping method. Recent advances in cell reprogramming technologies have opened up new ways to model neurological diseases by means of induced pluripotent stem cell (iPSC)-derived neurons obtained from patients. Many of the modeled diseases are associated with impaired synaptic connectivity [23] and experiments, as detailed below, conducted in patient-derived cultures, could shed light on some of the earliest synaptic dysregulations. In addition, newly developed CMOS HD-MEA systems for *in vivo* applications [24, 25, 26] hold the promise that the developed method will also be useful in a 3D context, and a first evaluation could include the application of our optimization and validation pipeline to *in vivo* HD-MEA data. This way, one could assess the effect of *in vivo* network characteristics (e.g., synchrony) on the connection mapping performance.

Next, we consider some limitations of the proposed approach and try to discuss possible improvements. In our experimental *in vitro* data, we achieved an approximately 50% success rate in finding and accepting the presynaptic unit of a target spine. The main potential reasons for rejection are that the presynaptic cell either *i*) was not captured by the selection of recording electrodes, *ii*) showed little inter-network burst activity, or *iii*) exhibited strongly correlated activity with other neurons. Point *i* can easily be addressed by performing multiple paired recordings from the same target spine, while sequentially changing the set of extracellular recording electrodes, so that neurons across the entire HD-MEA surface can be probed (e.g., see Figure 1A). Alternatively, next-generation CMOS HD-MEAs, featuring an increased number of simultaneously usable read-out channels (~20’000) [27, 28], will provide an even more comprehensive coverage of the network. The other two reasons for rejection are also important to consider. We selected target synapses based on the spines showing at least some individual activity in a pre-recording period, while there was also a significant number of spines that either remained silent or were active mostly during network bursting periods. Therefore, addressing points *ii & iii* may be particularly beneficial in order to maximize the number of identifiable connections. The required desynchronization of network activity could be achieved, for example, by applying temporarily desynchronizing stimuli or pharmacological treatments or by using different development stages of the cell cultures. Moreover, the activated states of *in vivo* neural activity may be naturally conducive to the identification of connections [29]. Finally, some of the assumptions underlying our method could be adjusted to more precisely fit different experimental conditions. For example, we here assume a certain log-normal distribution of release probabilities as reported before [20]. However, the user may wish to incorporate extra information about a specific network. Our analysis pipeline, therefore, allows for a convenient adjustment of the release probability distribution. Moreover, we assumed that spine Ca^2+^ transients can be approximated by convolution of the presynaptic spike train with a simple exponential decay kernel without a rising phase. Such simple assumptions have been shown to perform very well for spike deconvolution methods based on Ca^2+^ imaging data [18]. Nonetheless, the performance of our connection mapping method may be improved by explicitly modelling the effect of potential synaptic interactions and short-term plasticity on the evoked Ca^2+^ transients. However, as we excluded periods of strong network bursting, where these phenomena are most likely to occur, the extra computational costs may not necessarily justify a presumably small gain in performance. In the current implementation, we used Ca^2+^ indicators to readout postsynaptic spine signals. Alternatively, the same approach could be applied using next-generation voltage indicators [30] to provide an even higher temporal resolution, which might make the correlations between spike trains and spine signals more robust (provided that a signal-to-noise ratio comparable to Ca^2+^ can be achieved).

We envision that the proposed method will facilitate various studies of dendrite organization, function, and development. Here we highlight some of the possible investigations that exploit the unique set of provided information, including precise spike patterns of the presynaptic units and of large parts of the network. The spatial organization of synapses is critical for the input-output transformation of a neuron [1, 2, 3]. It is now possible to address questions such as: Are there any location dependencies concerning features of presynaptic spiking patterns (e.g., firing rate, “bursting” behavior, and more)? Furthermore, following the identification of the postsynaptic unit, it is possible to calculate the spike-transmission probability (STP) between pre- and postsynaptic neurons. How are STP values organized across the dendritic tree during spontaneous network activity? Exploiting the combined information of presynaptic spike times and evoked postsynaptic events, neurotransmission and its modulation by presynaptic short-term plasticity [2] processes could now be examined in detail at individual synapses during spontaneous presynaptic spike patterns. Moreover, combined long-term HD-MEA recordings with additional periodic spine imaging can be used to address important questions concerning activity-dependent synapse development, by assessing, e.g., structural spine plasticity or changes in neurotransmission. Another possible research direction could include the probing of synaptic interactions. Those are conventionally investigated by glutamate uncaging experiments [31, 19], where presynaptic spiking information is not available. Our method could potentially enable researchers to identify multiple adjacent synaptic inputs and, subsequently, allow to examine interactions during spontaneous activity. In summary, we developed a versatile experimental setup and analysis pipeline for the identification and interrogation of monosynaptic connections that complements existing approaches.

## 4 Methods and Materials

### 4.1 High-density Microelectrode Array (HD-MEA) system and electrode selection

A complementary-metal-oxide-semiconductor (CMOS) based HD-MEA featuring 26’400 electrodes was used [13]. This HD-MEA system has 1’024 reconfigurable readout channels that can be simultaneously recorded from at 20 kHz sampling frequency. We packaged the HD-MEA chip with epoxy (Epo-Tek 353ND, 353ND-T, Epoxy Technology Inc., Billerica, MA, United States) to cover the gold bond wires and the printed circuit boards (PCBs) before use. We used large plastic rings (diameter of 35 mm) for maintaining the neuronal cultures on the chips so as to enable access and sufficient scanning movement of the high-magnification dipping lenses (40x, 60x) of the upright confocal microscope. The electrodes at a pitch of 17.5 *μm* were covered with electro-deposited platinum black to decrease electrode impedance and improve the signal-to-noise characteristics.

The electrode configurations for simultaneous readout were selected in two ways: *i*) A brief prescan of all electrodes was performed, and the active electrodes were subsequently chosen. When there were more than 1024 active electrodes, we occasionally generated 2-3 sets of configurations. *ii*) Alternatively, we first identified the position of the target cell on the chip and then selected the surrounding electrodes.

### 4.2 Animal use and cell culture preparation

All experimental protocols involving animals were approved by the Basel-Stadt veterinary office according to Swiss federal laws on animal welfare, and were carried out in accordance with the approved guidelines. Prior to culturing cells, the HD-MEA was sterilized for 40 min in 70% ethanol and rinsed 3 times with de-ionized (DI) water. Next, the electrode array was treated with 20 *μ*L of 0.05% (v/v) poly(ethyleneimine) (Sigma-Aldrich) in borate buffer (Thermo Fisher Scientific, Waltham, MA, United States) at 8.5 pH, for 40 min at room temperature. This coating improves cell adhesion and makes the surface more hydrophilic. The chips were then rinsed three times with DI water to remove the remaining solution. Next, we added 8 *μ*L of 0.02 mg mL^*−*1^ laminin (Sigma-Aldrich) in Neurobasal medium (Gibco, Thermo Fisher Scientific) to support the growth and differentiation of the cells. The chips with laminin were incubated for 30 min at 37°C. Cortices of E-18 Wistar rat embryos were dissociated in trypsin with 0.25% EDTA (Gibco) and counted with a hemocytometer as described previously [32]. We then seeded 15’000 to 20’000 cells on top of the electrode arrays. Subsequently, the chips were incubated at 37°C for 30 min before adding 2 mL of plating medium. The plating medium stock solution consisted of 450 mL Neurobasal (Invitrogen, Carlsbad, CA, United States), 50 mL horse serum (HyClone, Thermo Fisher Scientific), 1.25 mL Glutamax (Invitrogen), and 10 mL B-27 (Invitrogen). After 3 days, 50% of the plating medium were replaced by a growth medium, with stock solutions consisting of 450 mL D-MEM (Invitrogen), 50 mL Horse Serum (HyClone), 1.25 mL Glutamax (Invitrogen), and 5 mL sodium pyruvate (Invitrogen). The procedure was repeated twice a week. The chips were kept inside an incubator at 37 °C and 5% CO_2_. All the experiments were conducted between days in vitro (DIVs) 18 and 21.

### 4.3 Fluorescence imaging of dendritic spine Ca^2+^ signals

For confocal fluorescence microscopy, we used a Nikon NiE upright microscope, equipped with Yokogawa W1 spinning disk scan head, an Andor iXon Ultra EMCCD camera (Oxford Instruments), and 40x/0.80 NA or 60x/1.00 NA water-objectives (Nikon). Cells were transduced with floxed jGCaMP7b (AAV1-syn-FLEX-jGCaMP7b-WPRE; Addgene #104493, MOI = 5×10^5^ vg) [14] and low-titer Cre (AAV9-hSyn-Cre-WPRE-hGH; Addgene #105553, MOI = 5×10^3^ vg) AAVs on DIV 7. A customized clamping device had parafilm M attached to the objective and the culture to avoid water evaporation during long recording sessions. Confocal image time series containing the target postsynaptic cell were acquired at 7-15 Hz, and large-scale electrical recordings of extracellular signals were simultaneously performed with the HD-MEA setup. A typical recording length was 2-3 min. Synchronization was achieved by sending TTL pulses from the camera to the field-programmable gate array (FPGA) controller of the HD-MEA system for post-recording alignment.

### 4.4 Processing of Ca^2+^ fluorescence traces

Fluorescence traces were initially extracted by averaging over the pixel values in the ROI for each frame. Next, the ΔF/F was computed by extracting the baseline value with a moving percentile function (25^th^ percentile, 100 data points as the moving stride). This also effectively detrended the signal. In order to isolate synaptically-evoked Ca^2+^ transients from the fluorescence traces, we removed the contamination by Ca^2+^ signals associated with backpropagating action potentials (bAP) of the imaged cell. To achieve this, a robust linear regression was performed using the dendritic shaft and spine traces and the contribution of the dendritic shaft (i.e., the bAP-component) was subtracted from the spine trace [15].

### 4.5 Processing of extracellular recordings

The extracellular recordings were first spike-sorted using SpyKING Circus 0.8.4 (width of templates=3 ms, spike threshold=6, cut-off frequencies for band-pass Butterworth filter=300 Hz/9500 Hz, spatial radius considered=210 *μ*m), followed by manual curation of each recording. Templates with a similar extracellular signature were deemed likely to be recorded from the same neuron and the spike time cross-correlograms were employed to support merging.

### 4.6 Burst detection based on global firing rate

To detect periods of strong synchronized network bursting, we analyzed variations in the population firing rate. As a first step, we computed individual unit spike rates. Next, hyperactive units were excluded to ease the computation of burst boundaries later in the procedure. To do so, we followed two strategies: i) to exclude units whose rates were beyond the 99.75^th^ percentile of the distribution of unit firing rates or ii) to exclude units whose rates were above 15 Hz.

The population rate of the clean units was computed in 10 ms bins. The 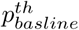 and 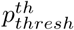 percentiles of these rates were assigned as the baseline and threshold rates, respectively (*f*_*baseline*_ and *f*_*thresh*_). Peaks in the population rate trace that satisfied the following conditions were then identified as bursts: peak rate was at least *f*_*thresh*_ and its prominence was at least *f*_*thresh*_. Prominence, a measure of how much a peak stands out relative to its neighbors, was computed as the height of the peak relative to its lowest local valley [33]. Additionally, a minimal inter-peak interval of *t*_*ipi*_ was imposed.

Individual burst boundaries were computed in the neighborhood of each peak using a smoothed version of the network rate trace. Smoothing was performed using locally weighted linear regression over a span of *n*_*span*_ data points. We found that smoothing yielded more robust estimates of burst boundaries. The start and end of each bursting period was defined as the first time point at which the smoothed rate crossed *f*_*baseline*_ to the left and right of the peak.

Edge cases, where data began or ended during a burst, were excluded. Further, a minimal interburst interval of *t*_*ibi*_ was imposed. If bursting periods were closer than *t*_*ibi*_, they were merged into a single larger burst.

The following parameter values were used in our analysis: *p*_*basline*_=85, *t*_*ipi*_=50 *ms, n*_*span*_=5, *t*_*ibi*_=75 *ms*. Depending on the degree of synchrony observed in the data, *p*_*thresh*_ was visually chosen for each data set. Lower values resulted in a less stringent definition of a bursting period, while higher values imposed more stringent conditions, resulting in much smaller periods being marked as bursts. Our choice of *p*_*thresh*_ typically lays between the 96^th^ and the 99^th^ percentile. Finally, we extended the detected bursting periods by 0.1 s before the burst onset and 0.5 s after the burst, in order to account for the slow dynamics of the Ca^2+^ signal.

### 4.7 Simulation of synaptically-evoked spine Ca^2+^ transients

We used a set of release probabilities following a log-normal distribution [20] (Figure 2B). We used the `numpy.random.lognormal` function to generate a set of log-normally distributed release probablities with the mean value of 2.5 and the standard deviation of 0.6 and scaled it by 0.01 to shift the values towards the desired range. Some very few values above 1 were replaced by random values between 0 to 1. We selected a background ROI close to the spine to measure the noise values and used the spine ROI as the signal region. In order to estimate the signal-to-noise ratio (SNR), the same number of pixels were extracted from the noise region and the signal region. The standard deviations of individual pixel traces were calculated, and we used the median values of the standard deviations of the signal and noise regions as the amplitudes of the simulated signal and noise, respectively. Then, the simulated spine response could be written as:

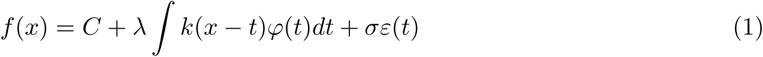

where *C* was the baseline of the fluorescence trace, *λ* denoted the amplitude of the pure signal (difference between the standard deviation of the measured signal and recording noise), *k*(*x* − *t*) was the instantaneous firing rate of the spike train, which was convolved with the exponential kernel *φ*(*t*) (decay *τ* = 0.5 *s*), and *σε*(*t*) indicated the recording noise term, as *σ* was the amplitude of the recording noise and *ε*(*t*) was white noise [34].

### 4.8 Binary classification

As described in the results section, we performed binary classification using the outcomes from correlation tests with and without the ground-truth units. A set of quantile thresholds ranged from the 0^th^ to 100^th^ quantile (Figure 4B) of the two distributions of GT-Rs and test-Rs. Note that in rare cases, the best-R of the correlation test with ground-truth unit was not from the ground-truth unit. For such an outcome, we classified it as FP if it passed the thresholds, or TN if it did not. Then the *F* score was calculated as follows:

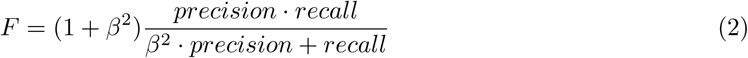

Here we used 0.5 for *β*, as precision weighs more than recall in the test. The optimal thresholds could be identified when the *F*-score reached its maximum.

## 5 Data availability

All the figures and results can be reproduced using the notebooks available on: https://github.com/starquakes/mea_spineca_mapping.git.

The extracellular recordings, imaging data, and curated spike sorting outputs have been converted to Neurodata Without Borders (NWB) [35, 12] using the `SpikeInterface` [17], `RoiExtractors` [36], and `nwb-conversion-tools` [37] software frameworks. The collection of NWB files is available on DANDI archive at: https://gui.dandiarchive.org/#/dandiset/000223.

## 6 Acknowledgements

This work was supported by the ERC Advanced Grant 694829 “neuroXscales”, the Swiss National Science Foundation project 205320-188910, the China Scholarship Council (X.X.), and the ETH Zurich Postdoctoral Fellowship 19-2 FEL-17 (A.P.B).

